# Contrasting effects of chronic lithium, haloperidol and olanzapine exposure on synaptic clusters in the rat prefrontal cortex

**DOI:** 10.1101/2020.04.14.033944

**Authors:** Els F. Halff, Marie-Caroline Cotel, Sridhar Natesan, Richard McQuade, Chris J. Ottley, Deepak P. Srivastiva, Oliver D. Howes, Anthony C. Vernon

## Abstract

The pathophysiology of the majority of neuropsychiatric disorders, including schizophrenia and mood disorders, involves synaptic dysfunction and/or loss, manifesting as lower levels of several presynaptic and postsynaptic marker proteins. Whether chronic exposure to antipsychotic drugs may contribute to this pattern of synaptic loss remains controversial. In contrast, the mood stabiliser lithium has shown to exhibit neurotrophic actions and is thought to enhance synapse formation. Whilst these data are not unequivocal, they suggest that antipsychotic drugs and lithium have contrasting effects on synapse density. We therefore investigated the effect of chronic exposure to lithium and to two different antipsychotics, haloperidol and olanzapine, on presynaptic Synaptic Vesicle glycoprotein 2A (SV2A) and postsynaptic Neuroligin (NLGN) clusters in the rat frontal cortex. Chronic exposure (28 days) to haloperidol (0.5 mg/kg/d) or olanzapine (7.5 mg/kg/d) had no effect on either SV2A or NLGN clusters and no overall effect on synaptic clusters. In contrast, chronic lithium exposure (2 mmol/L eq./d) significantly increased NLGN cluster density as compared to vehicle, but did not affect either SV2A or total synaptic clusters. These data are consistent with and extend our prior work, confirming no effect of either antipsychotics or lithium on SV2A clustering, but suggest contrasting effects of these drugs on the post-synapse. Although caution needs to be exerted when extrapolating results from animals to patients, these data provide clarity with regard to the effect of antipsychotics and lithium on synaptic markers, thus facilitating discrimination of drug from illness effects in human studies of synaptic pathology in psychiatric disorders.

## Introduction

Schizophrenia (SCZ) and Bipolar Disorder (BD) are psychiatric disorders that affect approximately 0.5% and 1% of the global population, respectively (WHO). For each of these disorders there is accumulating evidence that synaptic dysfunction is a hallmark of the pathophysiology of the disease. Neuroimaging and *post-mortem* studies of individuals with SCZ or BD show reduced brain volume and spine density, particularly in the prefrontal and anterior cingulate cortex and hippocampus relative to healthy controls^1–4^. These changes coincide with reduced protein and/or mRNA levels of various synaptic markers^5–11^. Whereas the majority of these data were derived from *post-mortem* human material, the recent development of the UCB-J PET tracers, which specifically interact with the presynaptic Synaptic Vesicle glycoprotein 2A (SV2A), has enabled visualisation of pre-synaptic terminal density in the living human brain^12–14^. SV2A is thought to play a role in neurotransmitter release and vesicle recycling^15–18^, and, due its localisation at both inhibitory and excitatory synapses throughout the brain, is considered as a proxy for overall synaptic density. Consistent with the aforementioned *post-mortem* evidence, studies using [^11^C]-UCB-J PET imaging provide evidence for a reduction in SV2A binding in chronic SCZ^19^. Whilst there is also evidence for reduced SV2A ligand binding in major depressive disorder^16^, data for BD is as yet lacking.

Our recent study in chronic SCZ patients also demonstrated that total SV2A protein levels and [^3^H]-UCB-J radioligand specific binding in the rat frontal cortex were unaffected by chronic exposure to two different antipsychotic drugs (APD)^19^. These data suggest that the reduced SV2A ligand binding observed in patients is related to SCZ pathophysiology rather than an effect of antipsychotic treatment. In contrast, there is no data concerning the effect of mood stabilising drugs used in the treatment of BD such as lithium (Li) on SV2A *per se*. Importantly, as SV2A is located in pre-synaptic terminals, analysis of SV2A alone does not fully address the question of whether exposure to either antipsychotic drugs or Li affects synaptic density or not. In this context, several studies investigating the effect of antipsychotic and mood-stabilising medication on the animal brain provide evidence to suggest that synaptic changes may occur upon administration of these drugs. These include altered spine density^20^, dendritic branching^21^, and altered expression of synaptic protein markers such as Synapsin, SNAP-25, Synaptophysin, and PSD-95^22–24^, however SV2A was not assessed in these studies. In earlier work we demonstrated contrasting effects of the antipsychotic drugs Haloperidol (HAL) and Olanzapine (OLZ) compared to the mood stabiliser lithium (Li) on rat cortical grey matter volume, showing an increase upon Li treatment but reduced cortical volume upon exposure to HAL and OLZ^25,26^. To gain further insight into the effect of psychotropic medication on synaptic density, we therefore investigated the effects of HAL, OLZ, and Li treatments on SV2A and Neuroligin (NLGN) clusters as global pre- and postsynaptic marker, respectively. We hypothesised that cluster count would be unaffected by chronic exposure to either HAL or OLZ, concurrent with our earlier data^19^, however we anticipated that chronic exposure to Li, which has been suggested to have neurotrophic and neuroprotective effects^21,27^, would lead to an increase in synaptic clusters. Our research provides one of the first investigations into the effect of these different psychotropic medications on SV2A clusters.

## Materials and Methods

### Animals

Animal experiments were carried out in accordance with the Home Office Animals (Scientific Procedures) Act (1986) and European Union (EU) Directive 2010/63/EU, with the approval of the local Animal Welfare and Ethical Review Body (AWERB) panel at King’s College London (KCL). Male Sprague-Dawley rats (Charles River UK Ltd, Margate, UK), initial body weight 220-270 gram (6-10 weeks of age) were housed four per cage in conventional plastic cages (38 × 59 × 24 cm, Tecniplast, UK) containing sawdust, paper sizzle nest and cardboard tunnels. Animals were maintained under a 12-hour light/dark cycle (07.00 lights on) with food and water available ad libitum. Room temperature and humidity were maintained at 21 ± 2°C and 55 ± 5%, respectively. Animals were habituated for a minimum of 7 days before experimental procedures.

### Experimental Design (animals)

Two separate batches of animals received continuous administration of psychotropic medication using osmotic minipumps (Alzet Model 2ML4; Alzet, Cupertino, CA, USA) for 28 days (equivalent to approximately 2.5 human years based on 11.8 rat days as equivalent to 1 human year^28^). Specifically, cohort 1 comprised animals of initial body weight 240-270 gram (9-10 weeks of age), treated with a common vehicle (β-hydroxypropylcyclodextrin, 20% wt/vol, acidified by ascorbic acid to pH 6; n=11), 0.5 mg/kg/day haloperidol (HAL; n=11), or 2 mmol/L equiv/kg/day lithium chloride (Li; n=10). Cohort 2 comprised animals of initial body weight 220-240 gram (6-7 weeks of age) treated with vehicle (n=4), or 7.5 mg/kg/day olanzapine (OLZ; n=12) (all chemicals from Sigma-Aldrich, Gillingham, UK). Blood plasma levels achieved using these doses and delivery by osmotic pumps are consistent with clinically comparable dopamine D2 receptor (D2R) occupancy and plasma levels^25,26,29^. With the exception of Li-treated animals, data reported in this study is derived from the same animals as those used in our previous study^19^.

Minipumps filled with drug or vehicle solutions were inserted subcutaneously on the back flank under isoflurane anaesthesia (5% induction, 1.5% maintenance delivered in an 80/20% medical air/oxygen mix). Weight was monitored daily in the week following surgery, and twice a week in the following weeks. Dyskinetic behaviour, *i.e.* vacuous chewing movements (VCMs) were measured at 2 and 4 weeks *post* surgery, during a 2-minute period following a 1-minute habituation phase outside the home cage. VCMs are stereotypical motor behaviours that commonly develop in animals treated with antipsychotics^30,31^ and are considered a proxy measure for tardive dyskinesia (TD) observed in patients undergoing chronic antipsychotic treatment. Animals undergoing Li treatment were given access to 0.9% saline instead of tap water to minimise hyponatremic properties of Li^32^.

### Post-mortem tissue handling and blood plasma analysis

After 28 days of administration, animals were terminally anaesthetised by injection of sodium pentobarbital (60 mg/kg, intraperitoneal) and culled by cardiac perfusion using heparinised (12.5 U/ml) ice-cold PBS. In order to prevent masking of synaptic target proteins and obtain a better quality of immunostaining^33,34^, animals were not perfusion-fixed. Following PBS perfusion, brains were extracted, hemisected and post-fixed overnight in 4% PFA at 4°C. Brain hemispheres were then washed once in PBS, incubated in PBS-buffered 30% sucrose solution for 48h at 4°C, and then snap-frozen on dry ice and stored at −70°C until further processing for immunostaining.

At termination a blood sample was collected for estimation of drug levels. HAL and OLZ levels were commercially measured using tandem mass spectrometry (Cyprotex, Macclesfield, UK); Li levels were measured on an Thermo Scientific X-Series 2 ICP-MS (Durham University, Durham, UK) optimised for sensitivity and minimal oxide interferences, using an external calibration curve of 10, 25, 50 and 100 ppb standards (Romil, Cambridge, UK). Good agreement was seen between both Li 6 and 7 isotopes indicating that spectral interferences were minimal.

### Fluorescence immunostaining

The left hemisphere of each animal was serially sectioned (30 µm-thick coronal sections, interval 1/12, 360 µm spacing between sections within a series) on a cryostat at −18°C and stored in tissue cryoprotection solution (25% glycerol, 30% ethylene glycol, 45% 1x PBS pH 7.4, 0.05% azide) at −20°C until further processing. Free-floating sections from each brain in each group were washed for 10 min in phosphate buffer (PB; 0.1M) and 2×10 min in PBS. For antigen retrieval sections were incubated for 10-15 mins in 10 mM sodium citrate (pH 6.2) at RT, followed by a 15 min incubation in pre-heated 10 mM sodium citrate (pH 6.2) in a water-bath at 78°C. Sections were then allowed to cool down to RT in the same solution while gently shaking for 30 min. Sections were then washed twice (2×5 min) in PBS supplemented with 0.05% Triton-X100 and incubated for 3-4h in blocking solution (10% Normal Goat Serum, 1.5% BSA, 0.3% Triton-X100 in PBS), followed by overnight incubation at 4°C with primary antibody diluted in blocking solution (1:1000 Rabbit-α-SV2A, Abcam ab32942, Cambridge, UK; 1:300 Mouse-α-Neuroligin1/2/3/4, Synaptic Systems 129211, Göttingen, Germany). Sections were then washed in PBS (3×10 min) and incubated for 2h in secondary antibody diluted in blocking solution (1:1000 Goat-α-mouse AlexaFluor488, abcam ab150113; 1:1000 Goat-α-rabbit AlexaFluor555, abcam ab150090) and washed again in PBS (4×10 min). Finally, sections were mounted on Superfrost Plus slides (ThermoFisher Scientific, UK), and air-dried at RT for 1h before coverslipping with mounting medium containing DAPI (Vectashield; Vector Laboratories, Peterborough, UK).

### Confocal image acquisition and analysis

Overview images of fluorescent staining in brain sections were acquired in automated batch mode on a VS120-L100-W slide scanner (Olympus, UK) and 4x lens objective. Confocal images for quantification of synaptic clusters were acquired at the Wohl Cellular Imaging Centre using an Inverted Spinning Disk confocal microscope (Nikon, Japan) and 60x oil immersion lens objective (NA 1.4). Images were 102.65 × 102.65 µm in size (512 × 512 pixels), acquired as a stack spanning 6-10 µm, at an interval of 0.3 µm. Image stacks were acquired from 4-5 consecutive sections containing the pre-specified brain regions, either the PFC, (Bregma +4.2 to +2.5mm, 6 stacks per section of which 3 in layer II/III and 3 in layer V) or the ACC (Bregma +2.3 to +0.0 mm, 4 stacks per section of which 2 in layer II/III and 2 in layer V). Synaptic clusters were analysed in ImageJ (*https://imagej.net/Welcome*), using a previously published macro^35^ that we adapted for our images and analysis requirements. In brief, the macro performs the following operations: 3 consecutive optical sections (equating 0.9 µm total thickness) within each image stack were manually selected based on quality of staining and contrast in the image and were then combined by maximum intensity projection. After background correction using a rolling ball with radius of 5 µm, the image was thresholded automatically using the “Moments Dark” algorithm. The thresholded image for each channel was used as a mask to measure cluster count, size and staining intensity within the clusters in the original image. Clusters smaller than 0.12 µm^2^ and larger than 5 µm^2^ (SV2A) or larger than 2.4 µm^2^ (NLGN) were excluded. Overlap was calculated by dilating clusters in the thresholded images of each channel by 1 pixel on all sides and overlaying both dilated masks. Overlapping clusters smaller than 0.12 µm^2^ (i.e. 3 pixels) were excluded.

### Statistical analysis

Statistical analyses were performed using Prism version 8 (GraphPad Software, La Jolla, California, USA). Samples sizes were based on prior studies^19,25,26^. Two animals (one VEH-treated and one OLZ-treated) were excluded from cluster analysis on the basis of sub-optimal saline perfusion of the brain, which negatively impacted on staining quality. Data were tested for statistical outliers based on ROUT test (0.5%) but no outliers were identified. Data were tested for normal distribution using the Shapiro-Wilk test. Scores for VCMs and immunostaining data were analysed by two-way ANOVA, assessing treatment, time, and treatment × time interaction (VCM), or treatment, region, and treatment × region interaction (immunostaining), with post-hoc Bonferroni’s multiple comparison test where appropriate. Linear correlation analysis was computed based on Pearson’s correlation coefficient (2-tailed). Researchers were blinded to the treatment groups during sample preparation, imaging and image analysis.

## Results

### Behavioural analysis and blood plasma levels

Following osmotic pump implantation, rats were monitored for weight loss or gain. All animals gained weight comparably with no significant differences between drug treatments and their respective control groups (Supplementary Figure S1A,B).

Administration of HAL, OLZ or Li by osmotic pump achieved plasma levels (mean ± SD) of 3.03 ± 0.94 ng/mL for HAL; 15.5 ± 5.54 ng/mL for OLZ and 0.3 ± 0.04 mmol/L for Li, respectively.

Vacuous chewing movements (VCMs) were measured at 2 and 4 weeks after the start of drug exposure. Rats exposed to HAL or OLZ, but not Li, showed a statistically significant increase in VCMs as compared to their respective control groups (Supplementary Figure S1C,D; C, main effect of treatment: F(2,29)=17.18, p<0.001, η_p_^2^=0.55; D, main effect of treatment: F(1,15)=11.33, p<0.01, η_p_^2^=0.60). There was no effect of time or treatment × time interaction on VCM count. There was also no correlation between HAL or OLZ plasma level and the number of VCMs (Supplementary Figure S1E,F), consistent with our prior work^25^.

### Effect of Haloperidol and Lithium treatment on synaptic markers

Guided by our previous observation of contrasting effects of HAL and Li on rat cortical volumes, as measured using MRI^26^, as well as the hypothesis that APDs do not affect SV2A cluster density^19^, whereas Li may show a neurotrophic effect, we investigated the effects of these two treatments on both pre- and postsynaptic staining in the rat prefrontal cortex (PFC) and anterior cingulate cortex (ACC) (Fig.1). These brain regions are most commonly associated with synaptic dysfunction in SCZ, BD and MDD, as well as in putative alterations in synaptic protein levels upon psychotropic medication. SV2A is present at presynaptic terminals of both excitatory and inhibitory synapses throughout the brain, therefore in order to also capture all postsynaptic structures, we chose an antibody recognising all subtypes of Neuroligin (i.e. Neuroligin 1-4, NLGN), which are present at excitatory and inhibitory synapses^36^, rather than the commonly used postsynaptic density (PSD) marker PSD-95, which is primarily present at excitatory synapses^37^.

**Figure 1.**
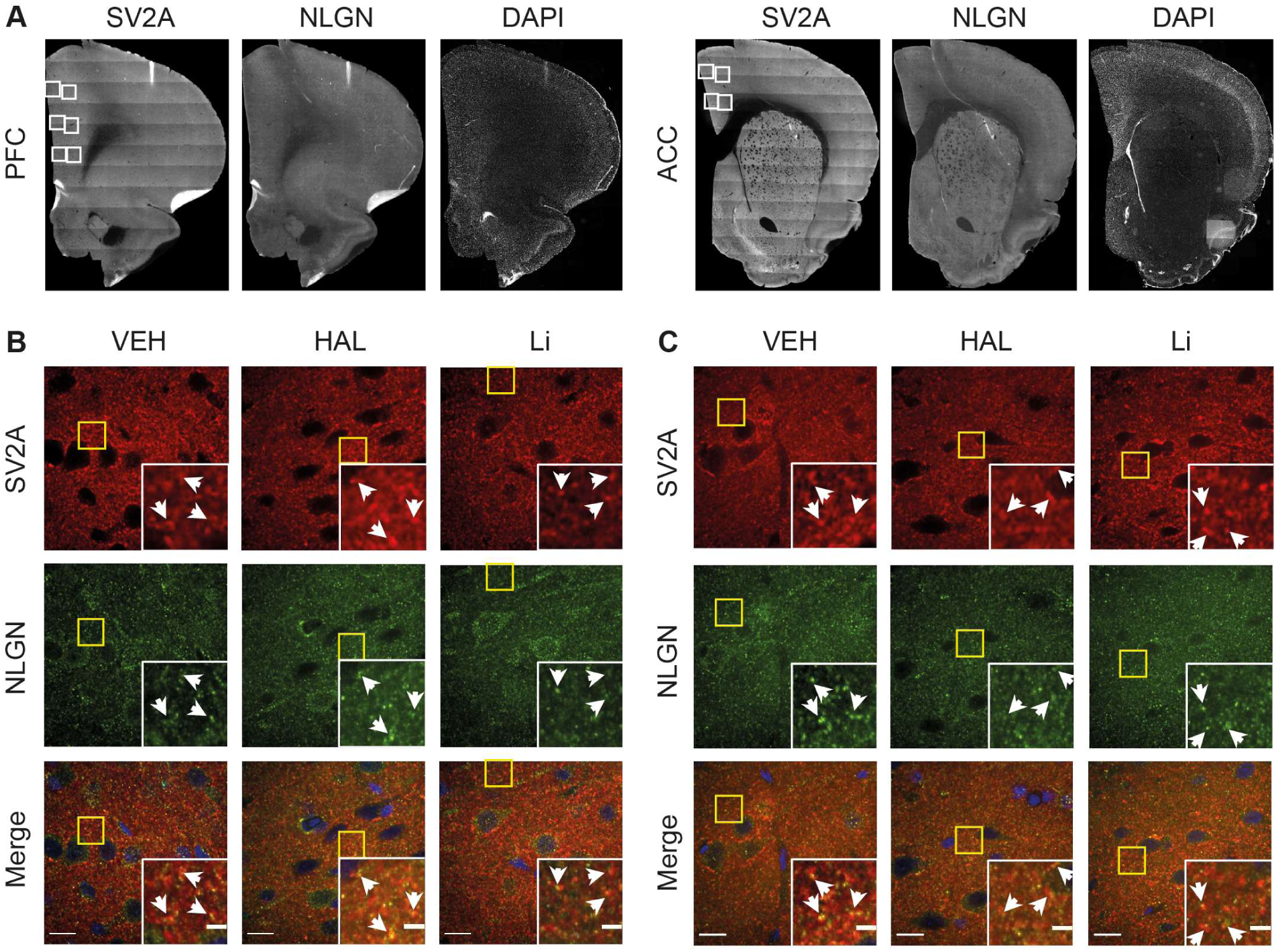
Confocal images of synaptic staining in rat PFC and ACC upon 28 days exposure to Haloperidol or Lithium. **A.** Fluorescent image (automatically stitched) of rat PFC (left) and ACC (right), stained with SV2A, NLGN, or DAPI as indicated. White squares indicate the positions where higher resolution confocal images (panel B,C) were acquired. **B**,**C.** Confocal images of the PFC (**B**) or ACC (**C**) showing synaptic staining with SV2A (top, red), NLGN (middle, green) or merged including DAPI (blue, bottom). Treatments are indicated at the top (VEH, Vehicle; HAL, 0.5 mg/kg/day Haloperidol; Li, 2 mM eq/kg/day Lithium). Yellow boxes indicate the position of the zoomed insets. Arrows indicate examples of co-localisation. Scalebar 15 µm (entire image) or 4 µm (zoomed insets).

Both SV2A and NLGN immunostaining were present throughout grey but not white matter (Fig.1A). At higher magnification, we found that SV2A immunostaining displayed clear punctate structures with high contrast in the pyramidal cell layers. Presynaptic SV2A structures were most prominent at perisomatic regions, but absent from the nucleus and soma (Fig.1B,C). NLGN staining revealed smaller clusters that were also present in the soma, but not in the nucleus (Fig.1B,C). We determined the number of pre- and postsynaptic clusters individually, as well as their overlap as a measure of synaptic density. Furthermore, we determined the total area of clusters as a measure for total synaptic terminal area, and the intensity of staining within clusters as a relative measure of SV2A and NLGN protein levels (Fig.2).

**Figure 2.**
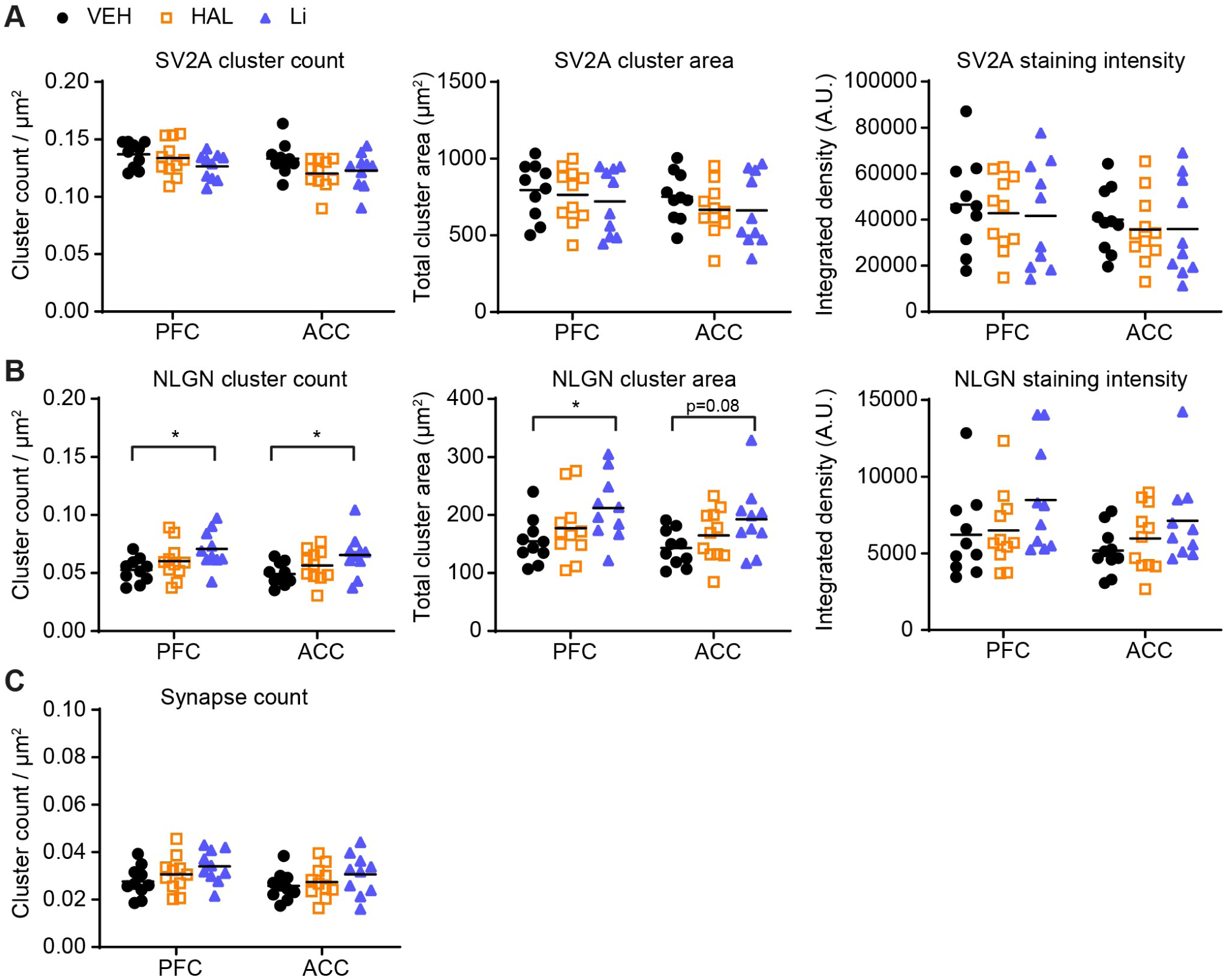
Quantification of SV2A and NLGN clusters upon 28 days exposure to Haloperidol or Lithium. **A-B.** Scatter plots of cluster count (left), cluster area (middle) and staining intensity (right) for rat PFC and ACC based on staining for SV2A (**A**), or NLGN (**B**). Treatments are indicated at the top (VEH, Vehicle, black circles; HAL, 0.5 mg/kg/day Haloperidol, orange squares; Li, 2 mM eq/kg/day Lithium, blue triangles). **C**. Scatter plot of cluster count of overlapping SV2A/NLGN clusters. Average value represents mean; two-way ANOVA with Bonferroni’s correction; *p<0.05.

Analysis of SV2A cluster number showed a main effect of region, but no significant effects of treatment or region × treatment interaction. Likewise, there were no effects of treatment or region × treatment interaction for the total area of SV2A clusters or total intensity inside clusters (Fig.2A, Table 1). Thus, our data suggests that chronic exposure to either HAL or Li does not affect SV2A-containing presynaptic terminals, and, by inference, presynaptic terminals *per se*.

**Table 1.** Statistical analysis (2-way ANOVA) of SV2A and NLGN staining in rat frontal cortex upon HAL and Li treatment

| Variable | Region | Treatment | Region x Treatment |
| --- | --- | --- | --- |
| SV2A cluster count | F(1,28)=10.40; **p<0.01**; $\eta_p^2=0.27$ | F(2,28)=2.072; ns<sup>1</sup> | F(2,28)=2.442; ns |
| SV2A cluster area | F(1,28)=7.570; **p<0.05**; $\eta_p^2=0.21$ | F(2,28)=0.5517; ns | F(2,28)=0.4628; ns |
| SV2A staining intensity | F(1,28)=7.354; **p<0.05**; $\eta_p^2=0.21$ | F(2,28)=0.2032; ns | F(2,28)=0.0268; ns |
| NLGN cluster count | F(1,28)=4.376; **p<0.05**; $\eta_p^2=0.14$ | F(2,28)=4.203; **p<0.05**; $\eta_p^2=0.64$ | F(2,28)=0.1007; ns |
| NLGN cluster area | F(1,28)=4.157; $p=0.051$ | F(2,28)=3.580; **p<0.05**; $\eta_p^2=0.57$ | F(2,28)=0.1245; ns |
| NLGN staining intensity | F(1,28)=8.478; **p<0.01**; $\eta_p^2=0.23$ | F(2,28)=1.986; ns | F(2,28)=0.5509; ns |
| Synaptic cluster count | F(1,28)=5.506; **p<0.05**; $\eta_p^2=0.16$ | F(2,28)=2.026; ns | F(2,28)=0.1408; ns |

<sup>1</sup>ns, not significant

Analysis of NLGN cluster number showed a significant effect of both region and treatment, but no region × treatment interaction (Fig.2B, Table 1). *Post-hoc* analysis revealed that the treatment effect was driven by a significant increase in NLGN cluster number in both PFC and ACC upon exposure to Li as compared to VEH-treated animals. Total NLGN cluster area also showed a significant increase upon Li treatment (Fig.2B, Table 1), although for the ACC this did not survive correction for multiple comparisons (actual p=0.08). There was no correlation between Li plasma level and NLGN cluster number or area (data not shown). Total NLGN staining intensity showed no significant effect of treatment (Fig.2B, Table 1). Thus, our data suggests that chronic Li, but not HAL exposure, induces an increase in postsynaptic cluster number and size.

Finally, we analysed the number of overlapping SV2A/NLGN clusters, as these would be representative of functional synapses. There was an overall effect of region on overlapping cluster count (Fig.2C, Table 1), suggesting an overall difference in synaptic density between PFC and ACC, but there were no statistically significant effects of either treatment or a treatment × region interaction.

### Effect of Olanzapine treatment on synaptic markers

To test whether the results obtained with HAL exposure generalises to other antipsychotics, we analysed synaptic clusters in a separate cohort of animals treated for 28 days with vehicle or 7.5 mg/kg/day Olanzapine (OLZ; Supplementary Figure S2A,B). Blood plasma levels of OLZ fell within the clinically relevant range. We found no effect of OLZ treatment on SV2A or NLGN cluster number, or overlapping synapse count (Supplementary Figure S2C, Table 2). Likewise, we found no effect on cluster area or staining intensity of either marker (Supplementary Figure S2D-E, Table 2). Thus, we show that OLZ, like HAL, does not affect pre- or postsynaptic clusters.

**Table 2.** Statistical analysis (2-way ANOVA) of SV2A and NLGN staining in rat frontal cortex upon OLZ treatment

| **Variable** | **Region** | **Treatment** | **Region x Treatment** |
| --- | --- | --- | --- |
| SV2A cluster count | F(1,14)=1.480; ns<sup>1</sup> | F(1,14)=0.1845; ns | F(1,14)=0.4974; ns |
| SV2A cluster area | F(1,14)=0.0268; ns | F(1,14)=1.222; ns | F(1,14)=0.0013; ns |
| SV2A staining intensity | F(1,14)=0.148; ns | F(1,14)=1.518; ns | F(1,14)=0.1523; ns |
| NLGN cluster count | F(1,14)=0.0605; ns | F(1,14)=0.7205; ns | F(1,14)=0.5051; ns |
| NLGN cluster area | F(1,14)=0.0063; ns | F(1,14)=0.9316; ns | F(1,14)=0.9005; ns |
| NLGN staining intensity | F(1,14)=0.1431; ns | F(1,14)=0.9287; ns | F(1,14)=2.076; ns |
| Synaptic cluster count | F(1,14)=0.235; ns | F(1,14)=0.8453; ns | F(1,14)=0.1283; ns |

<sup>1</sup>ns, not significant

## Discussion

### Summary and interpretation of results

Our study presented here investigates the effect of chronic exposure (28d) to the typical antipsychotic Haloperidol (HAL), the atypical antipsychotic Olanzapine (OLZ), and the mood stabiliser lithium (Li) on pre- and postsynaptic clusters using SV2A and NLGN as respective markers. In agreement with our hypothesis, we find that neither OLZ or HAL alter SV2A or NLGN synaptic cluster density, cluster area, or immunostaining intensity. These data confirm our recently published observations that neither HAL nor OLZ had any effect on either overall SV2A protein levels or [^3^H]-UCB-J specific binding in the rat PFC and ACC^19^. Our new data extends this prior work by demonstrating at higher resolution that, under the conditions tested, these APD also have no effect on synaptic clusters. Furthermore, we provide evidence for a contrasting effect of chronic Li exposure, which selectively enhances postsynaptic cluster count and area in both the PFC and ACC, without affecting SV2A-containing presynaptic terminal density or overall synapse count. The implications of these observations for clinical studies will be discussed further on.

This study is, to our knowledge, the first to investigate the effect of these psychotropic medications on SV2A clusters. Previous studies in rats have however investigated the impact of chronic exposure to APD on protein levels of Synaptophysin (SYN), a presynaptic vesicle protein that shows high correlation with SV2A distribution^12^. The majority of these studies found no statistically significant effects of either HAL or OLZ on SYN protein levels in the rat hippocampus and frontal cortex^23,38–40^; one study reported increased SYN levels in cortical lysates upon chronic administration of OLZ, however this study did not use clinically comparable dosing^41^. Supporting the majority of the rat studies, exposure of rhesus monkeys to clinically-comparable levels of HAL for >1 year found no change in SYN, either at DNA or protein levels, in cortical areas relevant to schizophrenia^42,43^. The effect of APD on postsynaptic marker expression has mostly been investigated at the mRNA level, with Homer1a and PSD-95 as the most commonly reported markers. For Homer1a mRNA levels in the cortex upon HAL and OLZ administration results are inconsistent, with some studies reporting upregulation and others downregulation^44–47^. These findings have yet to be confirmed at the protein level. For PSD-95 mRNA levels in the cortex findings were likewise inconsistent^45,47^, however no change was found for PSD-95 at the protein level in either striatum or frontal cortex^39,48^. Notably, both Homer1a and PSD-95 are primarily located at excitatory synapses therefore these results may not be reflective of overall synaptic protein levels.

Studies on the effects of chronic Li exposure on SYN reported no effect on either protein^40,41^ or mRNA levels^49^. On the other hand, Li exposure was shown to increase mRNA levels of Synapsins, a class of proteins that like SV2A and SYN is located on presynaptic vesicles and is involved in neurotransmitter release^22^, and is reduced in BD^40^. The latter study, however, did not see an effect of Li on Synapsin protein levels in the hippocampus. In cell culture, Li induced enhanced recruitment of both SYN and PSD95 to synapses^50^, indicative of a synaptogenic effect. Indeed, multiple studies confirm that Li treatment enhances dendritic branching and synapse formation, both *in vitro* and *in vivo*^21,51^. Thus, the neurotropic effect of Li may underlie the increase in postsynaptic clusters observed in our study.

Collectively, our data are consistent with these previous observations. Specifically, our results for SV2A parallel earlier studies on the effect of APDs and Li on SYN, whereas the effect of Li on increased NLGN clusters reflects the oft-reported neurotropic effect of Li. It should be taken into account however that direct comparisons are difficult to make as the literature is heterogeneous in terms of measures reported (protein *vs* mRNA), synaptic markers studied, regimens used to establish chronic exposure (daily injections, administration via food intake, or constant exposure via mini-pumps), length of exposure, as well as concentrations of drugs administered. Furthermore, due to slightly lower blood levels of Li in our samples, which fell just below the therapeutic range and earlier observed plasma levels^26^ the effect of Li treatment may not have been as robust in our study compared to other investigations, and the effect on overlapping synaptic cluster count upon Li exposure may be underestimated.

### Limitations

Limitations of our study should be noted. First, drug treatment was performed in otherwise healthy animals, therefore the interaction between drug exposure and illness was not assessed. Interestingly, OLZ partially rescued stress-induced reduction of SYN and PSD-95 protein levels in the rat prefrontal cortex^39^, as well as PCP-induced reduction of SYN and PSD-95 protein levels and neurite outgrowth in neuronal culture^52^, and PCP-induced reduction in spine density^53^, without affecting these measures in control condition. Future research should therefore include investigating the effect of APD and mood stabilising drugs on SV2A protein levels and synaptic cluster density in animal models relevant for the study of SCZ and mood disorder.

Second, the number of synapses detected is highly dependent on the synaptic marker, method of detection, and method of quantification used. It is not uncommon to see different numbers for pre- and postsynaptic markers, which will also be reflected by a lower number of overlapping (total) synapses as compared to the individual markers^54^ (Fig.2 and S2). Notably, our reported density of 0.1-0.2 SV2A clusters per µm^3^ is in the same order of magnitude as observations by Rasakham et al.^55^, who reported an approximate synaptic cluster density of 0.5 clusters per µm^3^ in the PFC of rats in a similar age range, using Synaptophysin and PSD95 immunostaining with fluorescence microscopy. At higher resolution, an electron microscopy study detected up to 6 synaptic clusters per µm^3^ in the rat PFC^56^, suggesting only a subset of synaptic clusters will be detected at the resolution of fluorescence microscopy. Thus, our study should be considered to reflect a percentage of change rather than a change in absolute cluster numbers.

Third, on the basis of our study we cannot exclude that APD exposure causes more subtle effects in synapse rearrangement that cannot be detected at the resolution of light microscopy. Electron microscopy studies in the rat PFC after up to 1 year of exposure to HAL showed a shift towards an increased number of inhibitory synapses without affecting total synapse number or synaptic vesicle density^57,58^. Furthermore, autoradiography studies from our lab revealed increased GABAA receptor binding in the rat ACC upon chronic HAL exposure^59^. Future studies combining SV2A staining with specific pre- and postsynaptic markers for inhibitory and excitatory synapses, such as vGAT/Gephyrin or vGlut/PSD95, respectively, may give more insight into the involvement of SV2A in regulating the excitatory/inhibitory balance (E/I balance) during disease progression and upon drug exposure.

In this context it is interesting to note that SV2A was found to be differentially associated with GABAergic and glutamatergic synapses depending on brain region and developmental stage^60,61^, and that an E/I imbalance has been associated with both SCZ and BD^62,63^. It therefore remains to be investigated whether a reduction of SV2A in psychiatric disorders truly represents a loss of synapses, or rather a disturbance in the E/I balance. Elucidating the functional impact of reduced SV2A levels on neurotransmitter release and synaptic signalling also remains a priority for future work.

### Implications for human studies

Our findings may have broader implications for clinical research into synaptic dysfunction in psychiatric disorder using both neuroimaging and *post-mortem* methods. SV2A is a synaptic protein of great interest due to the development of the UCB-J tracers, which have enabled *in vivo* imaging of presynaptic terminals in the living human brain^12,13,19^. One potential confounding factor in such human PET studies is whether psychotropic medication has an impact on the specific binding of the SV2A radioligand. To date, little is known about the effect of psychotropic mediation on SV2A, and *post-mortem* studies in experimental animals are thus essential for the correct interpretation of clinical PET studies^35^. Our data presented here provide evidence that APDs are unlikely to account for the reduced SV2A binding in SCZ patients as observed by PET^19^. More broadly, the lack of change in either SV2A, NLGN or synaptic (overlapping SV2A/NLGN) cluster density suggests APD treatment may be unrelated to changes in other synaptic markers found in human *post mortem* SCZ tissue, such as reduced SYN levels^8,64^. Whether SV2A protein levels are reduced in BD and SCZ *post-mortem* brain tissue has yet to be investigated, but could be expected to reflect the same changes as those observed with SYN.

The increase in NLGN cluster count and area upon Li treatment is consistent with the reported neurotrophic effects of this drug, although the mechanisms underlying this effect are as yet poorly understood^27^. At the resolution of our data we are unable to say whether the increase in NLGN clusters is due to an increase in the number of postsynaptic specialisations (such as increased spine density), or whether an increase in the size of pre-existing clusters due to postsynaptic protein accumulation^24^ leads to inclusion of clusters that were previously below detection level. It is interesting to note that we found a significant increase in NLGN clusters upon lithium treatment despite plasma levels falling just below the clinically relevant therapeutic range. It is tempting to speculate that the synaptogenic effects of Li do not underlie the improvement in the core BD symptoms of mania and depression, which are the main clinical measures to assess treatment efficacy in BD, but rather contribute to some therapeutic benefit of Li that is less well established such as reducing cognitive decline^65^. However, more work is needed to understand the exact implications of these results for the effect of treatment in BD.

In summary, our study shows that chronic APD exposure (up to 28 days) does not alter pre- or postsynaptic cluster density. In contrast, chronic Li exposure (up to 28 days) enhances postsynaptic clusters, although we saw no effect on the presynapse. Our data provide clarity with regard to the impact of psychotropic treatment on SV2A and synaptic density, which will aid in the interpretation of clinical studies. In particular, our findings reinforce the suggestion that reduced SV2A in SCZ represents a pathology intrinsic to the illness and not an adaptation to antipsychotic drug exposure^19^. Further work is however needed to understand how the apparent effects of Li on the post-synapse are related to its beneficial clinical effects on mood and cognition.

## Acknowledgements

We would like to thank George Chennell and Chen Liang of the Wohl Cellular Imaging Centre (King’s College London) for their valuable help with image acquiry.

ACV acknowledges financial support for this study from the Medical Research Council (New Investigator Research Grant (NIRG), MR/N025377/1). The work (at King’s College, London) was also supported by the Medical Research Council (MRC) Centre grant (MR/N026063/1). These funding sources had no further role in study design; in the collection, analysis and interpretation of the data; in the writing of the report; and in the decision to submit the paper for publication.

