## Supplemental Figure 1-2 for "Contrasting effects of chronic lithium, haloperidol and olanzapine exposure on synaptic clusters in the rat prefrontal cortex"

**Supplementary Figures and Figure Legends**

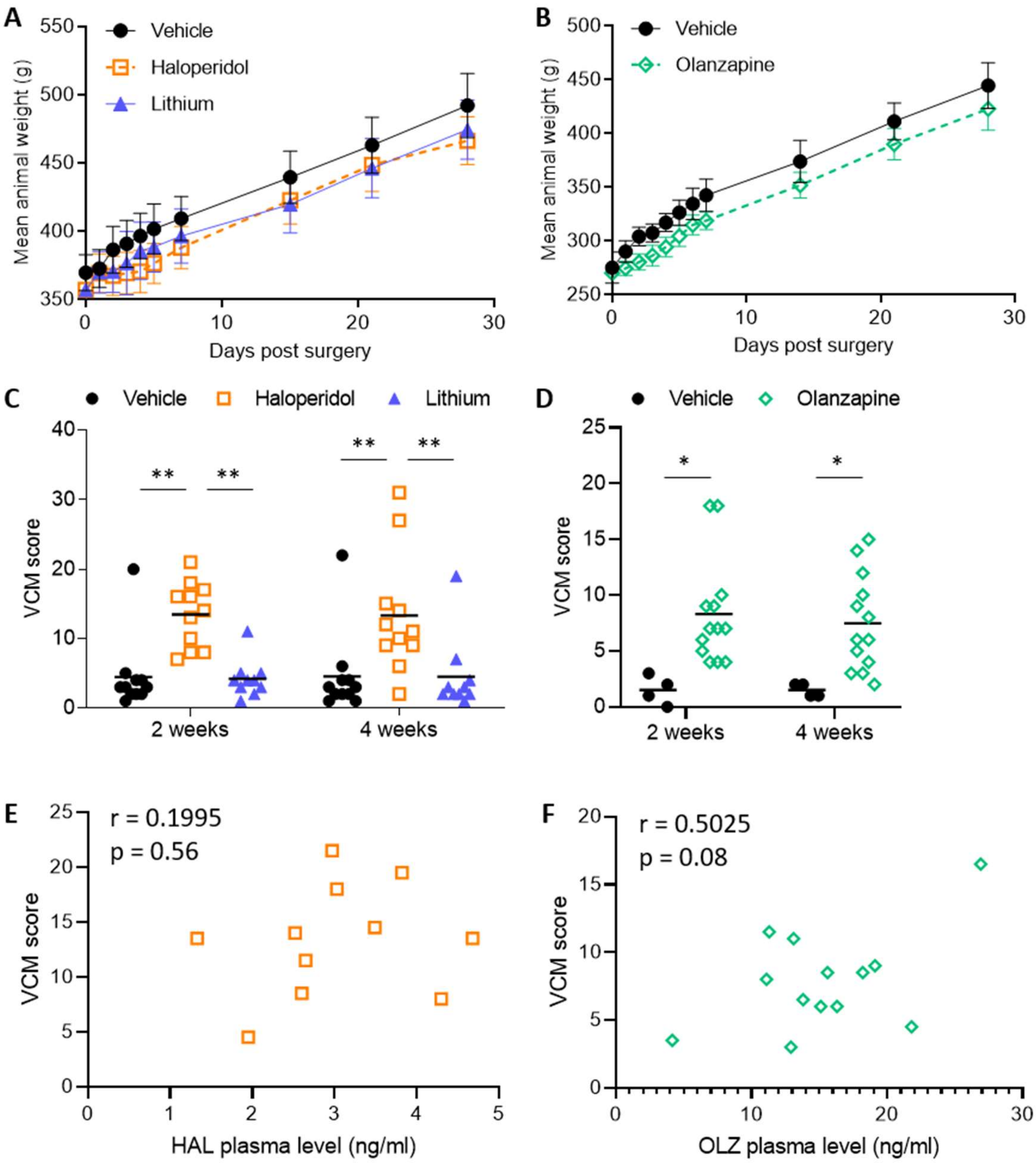


**Supplementary Figure S1. Weight monitoring and behavioural analysis of rats undergoing psychotropic treatment. A-B.** Weight recording of animals undergoing 28 days treatment with (**A**) Vehicle control, 0.5 mg/kg/day Haloperidol, or 2 mM eq/kg/day Lithium, and (**B**) Vehicle control, or 7.5 mg/kg/day Olanzapine. Average values represent the mean $\pm$ SEM. **C-D.** VCM scores measured at 2 or 4 weeks after the start of treatment (**C**: Vehicle control, 0.5 mg/kg/day Haloperidol, or 2 mM eq/kg/day Lithium; **D**: Vehicle control, or 7.5 mg/kg/day Olanzapine). Average values represent the mean; two-way ANOVA with Bonferroni's *post hoc* correction; *$p<0.05$, **$p<0.01$. **E-F.** Scatter plots of antipsychotic drug plasma levels *versus* VCM scores of animals exposed to Haloperidol (**E**) or Olanzapine (**F**). VCM scores represent the average score between week 2 and week 4 for each individual animal; Pearson's correlation test.

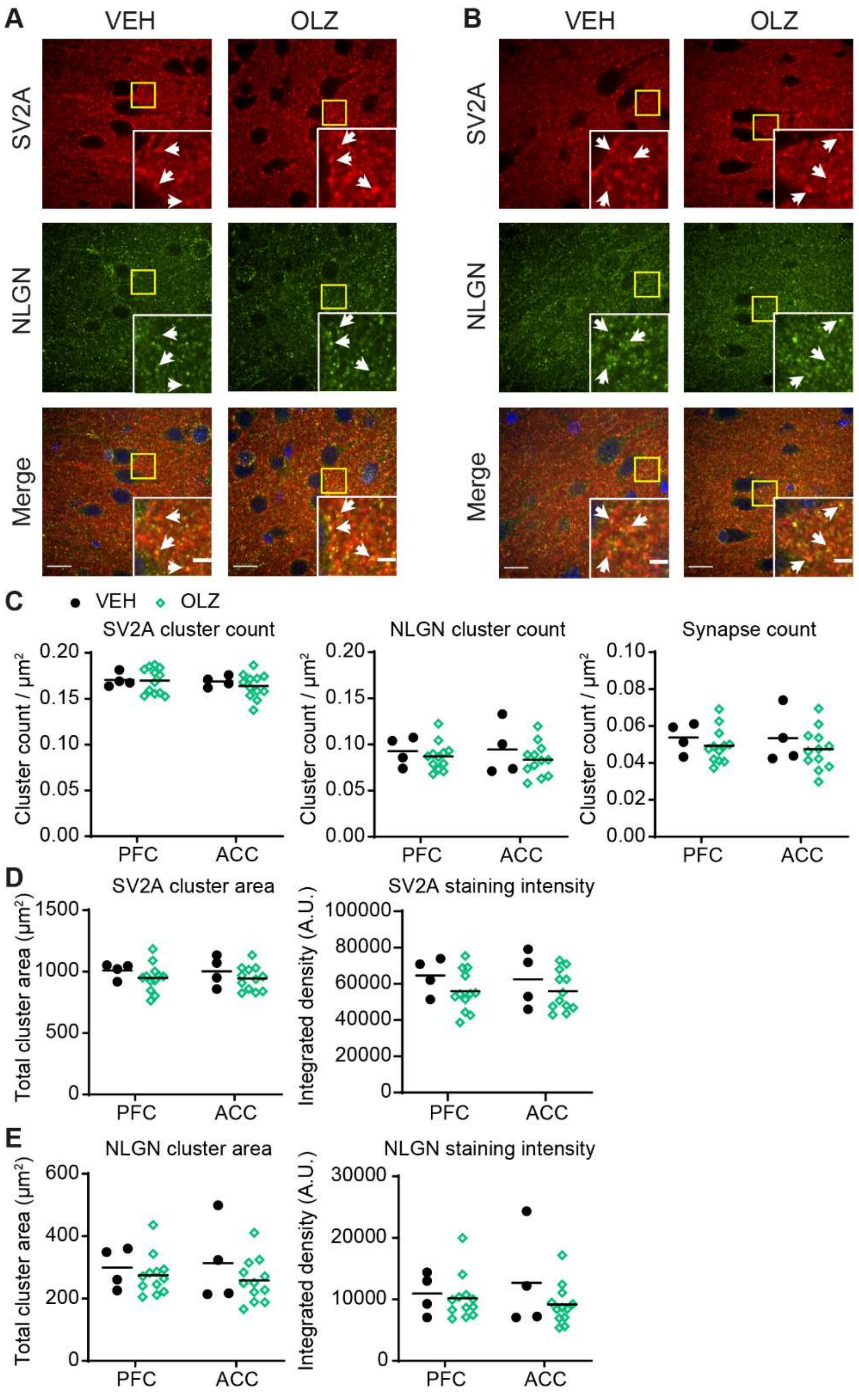


**Supplementary Figure S2. Synaptic staining and cluster analysis in the frontal cortex of rats upon 28 days exposure to Olanzapine. A-B.** Confocal images of the PFC (**A**) or ACC (**B**) showing synaptic staining with SV2A (top, red), NLGN (middle, green) or merged including DAPI (blue, bottom). Treatments are indicated at the top (VEH, Vehicle; OLZ, 7.5 mg/kg/day Olanzapine). Yellow boxes indicate the position of the zoomed insets. Arrows indicate examples of co-localisation. Scalebar 15 µm (entire image) or 4 µm (zoomed insets). **C.** Scatter plot of cluster count for rat PFC and ACC based on staining for SV2A (left), NLGN (middle), or SV2A/NLGN overlap (right). Treatments are indicated at the top (VEH, Vehicle, black circles; OLZ, 7.5 mg/kg/day Olanzapine, green diamonds). **D-E.** Scatter plots of cluster area (left) and staining intensity (right) for rat PFC and ACC based on staining for SV2A (**D**), or NLGN (**E**).

Average value represents mean; two-way ANOVA with Bonferroni's correction.
